# A retinotopic code structures the interaction between perception and memory systems

**DOI:** 10.1101/2023.05.15.540807

**Authors:** Adam Steel, Edward H. Silson, Brenda D. Garcia, Caroline E. Robertson

## Abstract

Conventional views of brain organization suggest that the cortical apex processes internally-oriented information using an abstract, amodal neural code. Yet, recent reports have described the presence of retinotopic coding at the cortical apex, including the default mode network. What is the functional role of retinotopic coding atop the cortical hierarchy? Here, we report that retinotopic coding structures interactions between internally-oriented (mnemonic) and externally-oriented (perceptual) brain areas. Using fMRI, we observed robust, inverted (negative) retinotopic coding in category-selective memory areas at the cortical apex, which is functionally linked to the classic (positive) retinotopic coding in category-selective perceptual areas in high-level visual cortex. Specifically, these functionally-linked retinotopic populations in mnemonic and perceptual areas exhibit spatially-specific opponent responses during both bottom-up perception and top-down recall, suggesting that these areas are interlocked in a mutually-inhibitory dynamic. Together, these results show that retinotopic coding structures interactions between perceptual and mnemonic neural systems, thereby scaffolding their dynamic interaction.

## Introduction

Understanding how mnemonic and sensory representations functionally interface in the brain while avoiding interference is a central puzzle in neuroscience^1–5^. Previous work has shown that representational interference can be reduced through operations like orthogonalization^1^, pattern separation^6^, and areal segregation^5^. However, the principles that preserve interaction across mnemonic and perceptual populations are less clear.

This puzzle is particularly perplexing because classic models of brain organization generally assume that perceptual and mnemonic cortical areas do not share neural coding principles. For example, the neural code that structures visual information processing in the brain – retinotopy – is not thought to be shared by mnemonic cortex. Instead, it is thought that retinotopic coding is replaced by abstract, amodal coding as information propagates through the visual hierarchy^7–10^ towards memory structures at the cortical apex^11–14^. How can visual and mnemonic information interact effectively in the brain if they are represented using fundamentally different neural codes?

Recent work has suggested that even high-level cortical areas, including the default mode network, exhibit retinotopic coding: they contain visually evoked population receptive fields (pRFs) with inverted response amplitudes^15,16^. But the functional relevance of this retinotopic coding at the cortical apex remains unclear. Here, we propose that the retinotopic code spanning levels of the cortical hierarchy links functionally-coupled mnemonic and perceptual areas of the brain, structuring their interaction. We investigated this proposal in three fMRI experiments.

## Results

We began by determining whether retinotopic coding was present in mnemonic and perceptual regions spanning levels of the cortical hierarchy. In Exp. 1, participants (N=17) underwent visual population receptive field (pRF) mapping^17,18^ **(Fig. 1A)** with advanced multi-echo fMRI^19,20^ to maximize signal-to-noise and pRF model fitting in anterior regions of the temporal and parietal lobes^20^. As expected, an initial group-level whole-brain analysis revealed robust, positive retinotopic responses extending from early to high-level visual areas (positive visually-evoked pRFs, +pRFs) **(Fig. 1B)**. We also observed robust and reliable pRFs beyond the anterior extent of known retinotopic maps^21^ in anterior ventral temporal and lateral parietal cortex, regions of the cortical apex historically considered amodal^12,13^ **(Fig. 1B).** In striking contrast with classically-defined visual areas, a large portion of these anterior voxels had pRFs with an inverted visual response (i.e., negative visually-evoked pRFs, -pRFs^15,16^). For brevity, we refer to voxels with positive or negative pRF response amplitudes as +/− pRFs, respectively. This agrees with prior studies that also observed -pRFs in the brain^15,16^, but the functional relevance of this inverted retinotopic code is unknown^10^.

**Figure 1.**
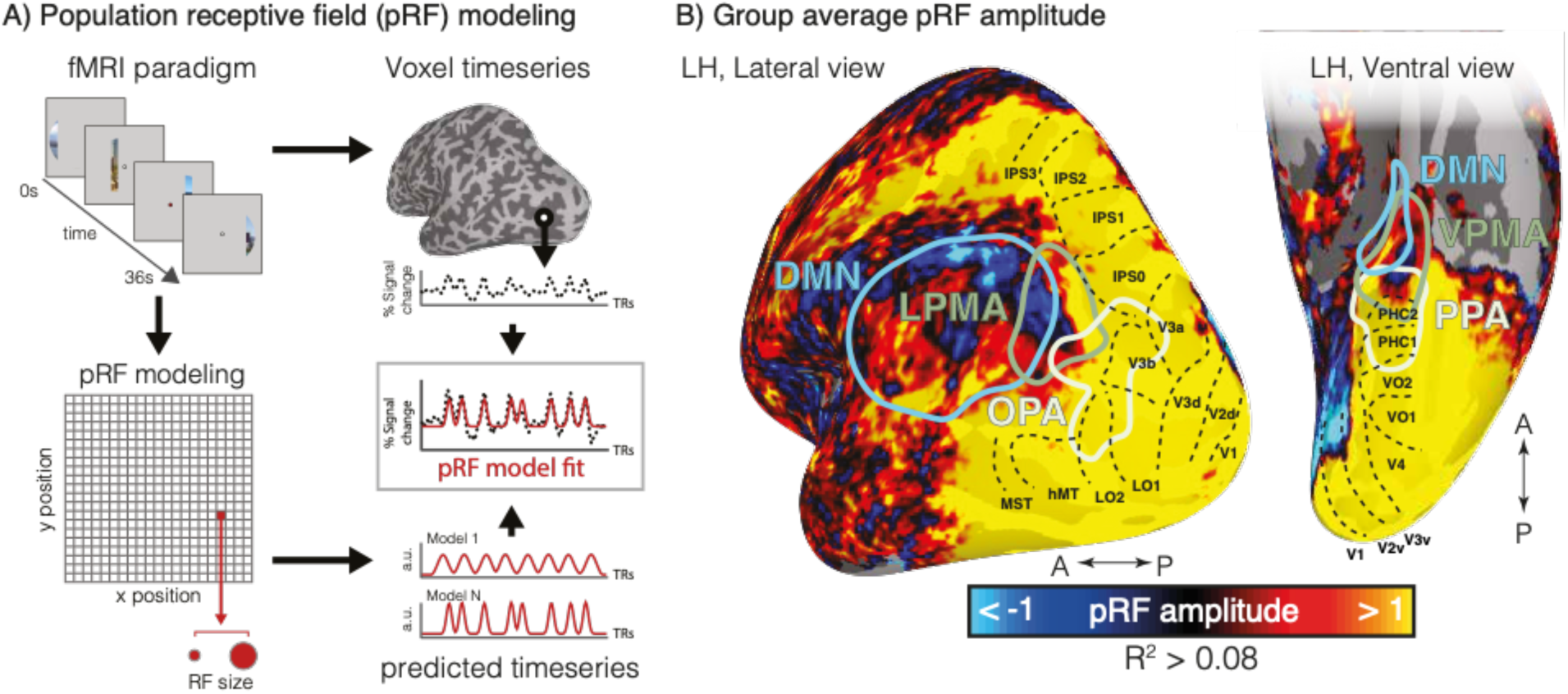
Population receptive field (pRF) mapping reveals retinotopic coding throughout posterior cortex, comprised of both positive and negative amplitude pRFs. **A.** PRF paradigm and modeling. In Exp. 1, participants underwent pRF mapping. Participants viewed visual scenes through a bar aperture that gradually traversed the visual field. Each visual field traversal lasted 36 seconds (18, 2s positions), and the bar made eight traversals per run. The direction of motion varied between traversals. To estimate the pRF for each voxel, a synthetic timeseries is generated for 400 visual field locations (200 x and y positions), and 100 sizes (sigma). This results in 4 million possible timeseries that are fit to each voxel’s activity. The fit results in four parameters describing each receptive field: x, y, sigma, and amplitude. **B.** Negative amplitude pRFs fall anterior to cortex typically considered visual (beyond known retinotopic maps). Group-average (n=17) pRF amplitude map (thresholded at R^2^>0.08) is shown on partially inflated representations of the left hemisphere, alongside regions-of-interest: scene perception areas (OPA, PPA), place memory areas (LPMA, VPMA) (localized in an independent group of participants^23^), and the default mode network ^24^. The known retinotopic maps in posterior cortex (black dotted outlines^21^) contain exclusively positive amplitude pRFs (hot colors), as visual stimulation evokes positive retinotopically specific BOLD responses. Negative amplitude pRFs (cool colors), where visual stimulation evokes a negative, spatially specific BOLD response, arise anterior to these retinotopic maps in the place memory areas and default mode network (see Fig. 2A for example time series from a representative subject).

At the group-level, -pRFs only appeared beyond the anterior extent of known retinotopic maps^21^, in areas associated with the cortical apex, consistent with prior reports^15,16^ **(Fig. 1B)**. This topographic profile of pRF amplitude was also evident at the single participant level, with robust pRF responses present throughout posterior cerebral cortex and -pRFs emerging at the anterior edge of high-level visual cortex (**Fig. S1**). In general, these anterior pRFs were not arranged topographically on the cortical surface in a manner that recapitulates the layout of the retina (i.e., did not constitute ‘retinotopic maps’). However, these -pRFs were robust and reliable within individuals: a half-split analysis (see Methods) confirmed that, for all ROIs, greater than 78% of pRFs had consistent signed amplitude (i.e., positive or negative), and these pRF amplitudes (all R>0.1, p<0.001) and visual field coverage maps (all dice-coefficients > 0.12, p<0.04, D>0.54) were all highly reproducible across splits. Importantly, these anterior -pRF populations on the ventral and lateral surfaces are distinct from the population of -pRFs observed on the medial wall in peripheral early visual areas, which arise from stimulation outside the pRF’s receptive field (i.e., surround suppression) ^16,22^.

If the inverted retinotopic code at the cortical apex serves a functional role of structuring interactions with perceptual areas, we hypothesized that -pRFs would concentrate in mnemonic, as compared with perceptual functional areas. Consistent with this prediction, at the group level we observed that -pRFs tended to fall within swaths of cortex that selectively responded during top-down recall of familiar places (place memory areas, PMAs^23^) on the lateral and ventral surfaces (lPMA and vPMA, respectively) **(Fig. 1B)**. These two mnemonic areas each lie immediately anterior to one of the scene perception areas^26^ (SPAs) of the human brain (occipital place area, OPA^25,26^, and parahippocampal place area, PPA ^27^). In contrast to the PMAs, the SPAs tended to contain +pRFs.

We confirmed this observation in individual participants by calculating the percentage of +/− pRFs within individually localized SPAs and PMAs (lateral: OPA, LPMA; ventral: PPA, VPMA) (**Fig. 2A**, see Methods) to better understand their nuanced functional topography^23^ **(Fig. 2A, S1-S3).** This analysis confirmed that, unlike the perceptual SPAs, which almost exclusively contained +pRFs, the mnemonic PMAs contained a significant percentage of -pRFs **(Fig. 2B)** (two-way rmANOVA, main effect of ROI (F(1, 16)=85.83,p<0.0001; ROI:Hemisphere interaction (F(1, 16)=11.24, p=0.004); post-hoc tests: lh: OPA v LPMA t(16)=10.74, p_corr_ =0.000002, D=3.43; rh: OPA v LPMA t(16)=5.97, p_corr_ =0.00002, D=1.89). On the ventral surface, the effect of greater -pRFs in the PMAs vs. SPAs was generally stronger in the left compared to the right hemisphere (two-way rmANOVA, main effect of ROI (F(1, 16)=10.01,p=0.006; no ROI:Hemisphere interaction (F(1, 16)=3.12, p=0.09; lh: PPA v VPMA t(16)=2.62, p_corr_=0.02, D=0.94; rh: PPA v VPMA t(16)=2.57, p_corr_=0.04, D=0.90). Notably, in many subjects, the PMAs contained both -pRF and +pRF subpopulations. Both -/+ pRFs within each memory area showed overall positive activation during a memory recall task (t-test versus zero: -LPMA vs +LPMA t(16)=17.67, p=6.35-12, D=4.28; -VPMA vs +VPMA t(11)=12.43, p=5.89-9, D=2.31; -LPMA vs +LPMA: (t(16)=0.44, p=0.65, D=0.04; -VPMA vs +VPMA: t(11)=0.26, p=0.79, D=0.07; **Fig. S6**). This suggests the +/− pRF subpopulations in the PMAs are comparably engaged in memory recall, although they show opposite response profiles to visual stimulation. Importantly, these results of a higher proportion of -pRFs in the PMAs vs. the SPAs held across a wide range of pRF R^2^ thresholds (R^2^ > 0.05 to R^2^> 0.20). Together, these results show that -pRFs are disproportionately represented in category-selective mnemonic areas (PMAs), as compared with their perceptual counterparts (SPAs).

**Figure 2.**
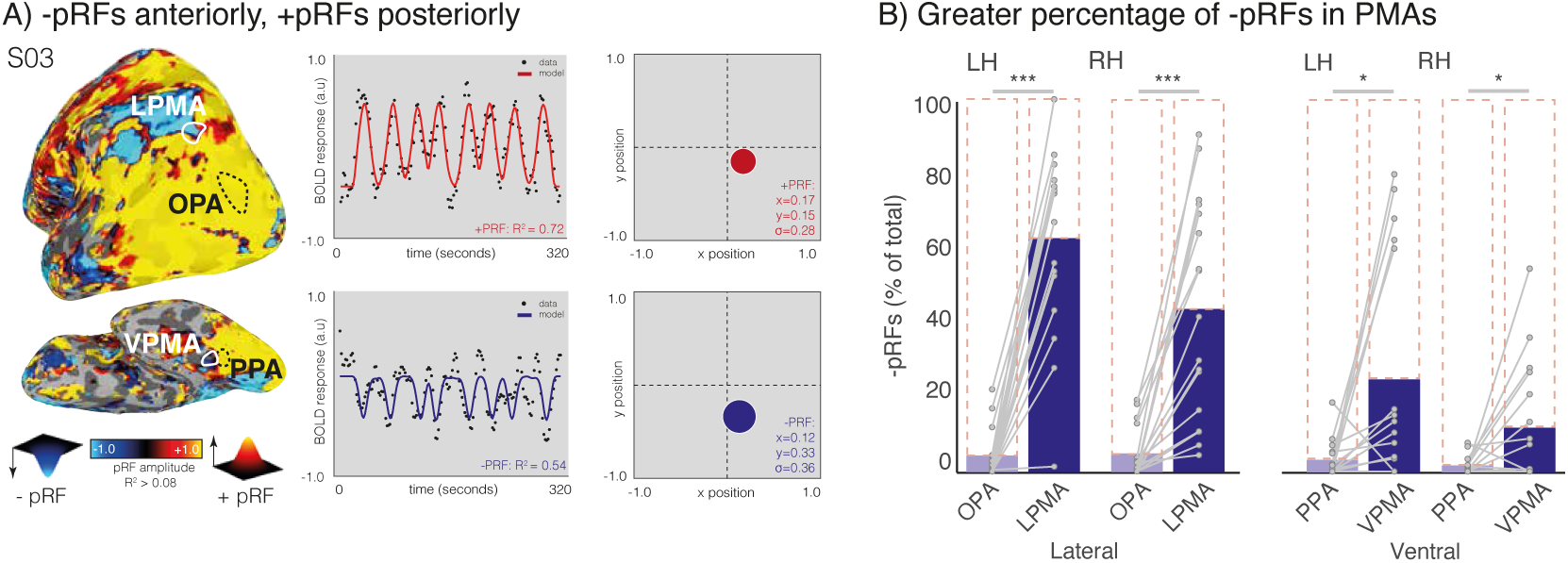
Transition to mnemonic cortex is marked by the appearance of negative pRFs. **A**. PRF modeling reveals posterior-anterior inversion of pRF amplitude in individual participants. Left. PRF amplitude for a representative participant overlaid onto a lateral view of the left hemisphere (thresholded at R^2^ > 0.15; see **Fig. S1** for example ventral and lateral surface pRF amplitude maps from all participants) and **Fig. S3** for amplitude maps with default mode parcellation overlaid. Posterior visual cortex is dominated by positive amplitude pRFs (hot colors), while cortex anterior to regions classically considered visual exhibits a high concentration of negative amplitude pRFs (cold colors). This individual’s lateral scene perception area (OPA) and place memory area (LPMA) are shown in white. Both the SPAs and PMAs contain pRFs **(Fig. S2)**. Right. Timeseries, model fits, and reconstructed pRFs for two surface vertices in this subject. (top) Example prototypical positive amplitude pRF from the lateral scene perception area (OPA). (bottom) Example negative amplitude population receptive field from the lateral place memory area (LPMA). **B.** Memory areas (PMAs) contain a larger percentage of negative pRFs compared to perceptual areas (SPAs). Blue bars depict percent of negative pRFs from individually localized SPAs and PMAs compared to total pRFs in the area (dotted outline). On the ventral and lateral surface, SPAs are dominated by positive pRFs, whereas a transition from positive to negative pRFs is evident within PMAs. Individual participant data points overlaid and connected in grey. *: p<0.05, ***: p<0.001, ns: non-significant.

Interestingly, both +pRFs and -pRFs in the PMAs tended to be smaller than +pRFs in the SPAs on both surfaces (Lateral surface: Main effect of ROI (F(2, 32)=10.27, p=0.0003; OPA vs +pRFs in LPMA: t(16)=5.61, Pcorr=0.0001, D=1.83; OPA vs -pRFs in LPMA: t(16)=2.59, p_corr_ =0.01, D=1.31; Ventral surface: F(2, 22)=15.46 p=0.000006; PPA v +VPMA: t(11)=2.83, p_corr_ =0.03, D=0.99; PPA v -VPMA(t(11)=5.43, p_corr_ =0.00006, D=1.85). PRF sizes were not significantly different between +/− pRFs in LPMA (t(16)=1.52, p_corr_ =0.14, D=0.48), but on the ventral surface +pRFs in VPMA were significantly larger than -pRFs in VPMA (t(11)=2.87, p_corr_ =0.03, D=1.06). This result of smaller pRFs in the PMAs compared with the SPAs is particularly notable, as it runs counter to the typical pattern of pRFs becoming larger moving anteriorly from early visual areas towards higher-level brain regions^8,17,28,29^. Although how the additional spatial precision manifests in these anterior areas is not clear, this result suggests that the PMAs could potentially represent highly specific information even compared to their perceptual counterparts **(Fig. 3)**.

**Figure 3.**
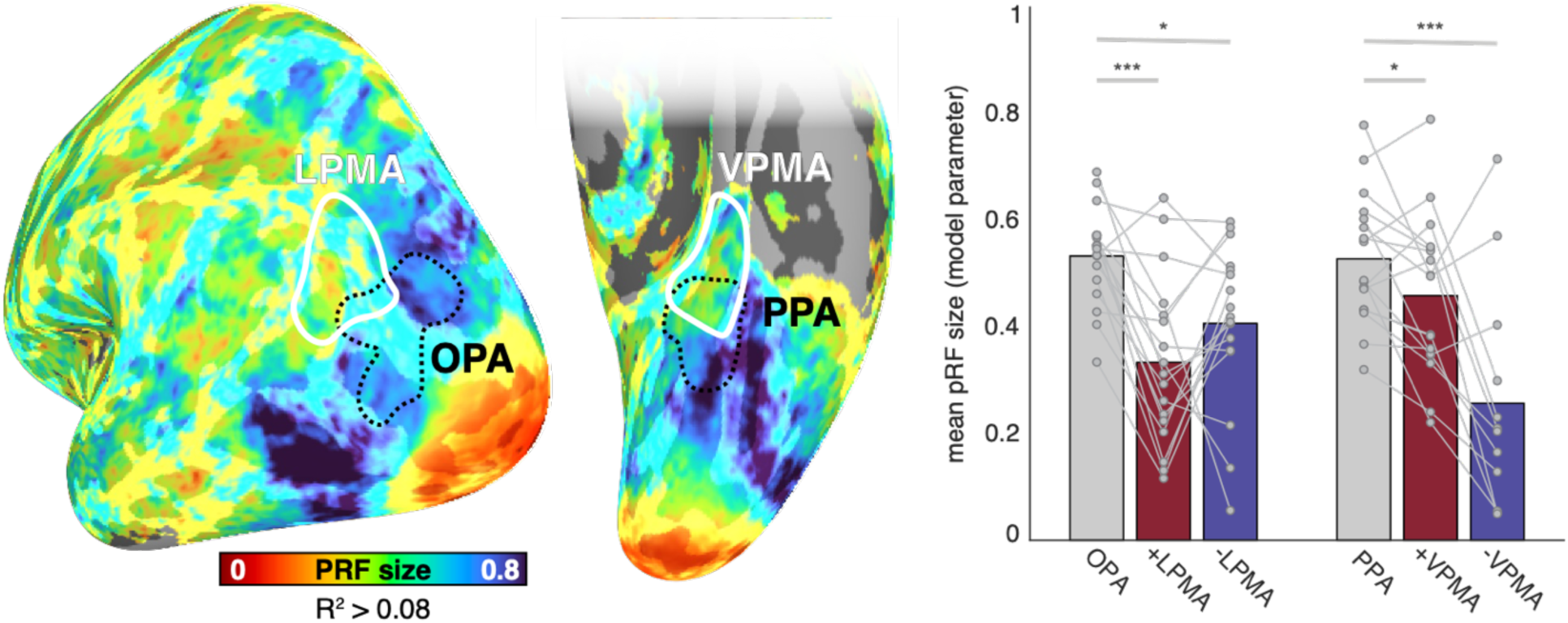
Memory areas contain smaller pRFs compared to their paired perceptual areas. **Left**. Group average pRF size with memory areas and perception areas overlaid. Nodes are thresholded at R^2^ > 0.08. **Right.** Bars represent the mean pRF size for +pRFs in SPAs (OPA, PPA) and +/− pRFs in PMAs (LPMA, VPMA). Individual data points are shown for each participant. Across both surfaces, pRFs were significantly smaller on average in the PMAs than their perceptual counterparts.

The enriched concentration of -pRFs in mnemonic (PMA) as compared with perceptual (SPA) areas led to a specific hypothesis regarding their functional role. Specifically, the PMAs are thought to act as a bridge between the perceptual SPAs and the spatio-mnemonic system in the medial temporal lobe^23^, but the format for sharing information between these systems is not clear. We hypothesized that retinotopic coding could serve as a shared substrate to scaffold the interaction between perceptual and mnemonic systems. If this proposal is correct, we should observe evidence for a functional interaction between these areas that depends on retinotopic position. We explored three tests of this hypothesis: 1) do paired +/− pRFs represent similar portions of the visual field (Exp. 1)?, 2) do the +/− pRFs identified during visual stimulation maintain their opponent activation profiles during top-down memory recall (Exp., 2)?, and 3) is the functional link between +/− pRFs recapitulated when viewing familiar scene images (Exp. 3)?

Next, we reasoned that, -pRFs within mnemonic areas should exhibit the same differential visual field biases as their perceptual counterparts^18^ if the retinotopic code scaffolds communication between perceptual and mnemonic systems. In particular, visual field representations in OPA and PPA are biased towards the lower and upper visual fields, respectively. Do -/+ pRFs in their respective PMAs share these biases? Using pRF data (Experiment 1), we calculated visual field coverage estimates for each pRF population (+pRFs in SPAs, +pRFs and - pRFs in PMAs). Consistent with prior reports, we found that OPA and PPA showed lower and upper field biases, respectively (Elevation bias OPA vs PPA: t(16)=2.42, p=0.02, D=0.98) **(Fig. 4A)**. Critically, we found that the visual field representations of the -pRFs in memory areas closely matched their paired perceptual counterparts **(Fig. 4B)**. Specifically, -pRFs in LPMA were biased towards the lower visual field (matching OPA), whereas -pRFs in VPMA were biased towards the upper visual field (matching PPA) (Elevation bias -LPMA vs -VPMA: t(11)=4.53, p=0.0008, D=0.94). In contrast, the +pRF populations in LPMA/VPMA did not show biases that corresponded to their perceptual counterparts: +pRFs in LPMA were biased towards the upper visual field, the opposite of OPA, and +pRFs in VPMA showed no clear elevation bias (Elevation bias +LPMA vs +VPMA: t(16)=2.78, p=0.01, D=0.65). These data show that the -pRFs in the PMAs represent similar visual field locations as their paired SPAs, and, based on the known functional interaction between the paired PMAs and SPAs during scene processing^23^, we suggest that the visual field representation may be, in part, inherited from feedforward connections originating in the SPAs.

**Figure 4.**
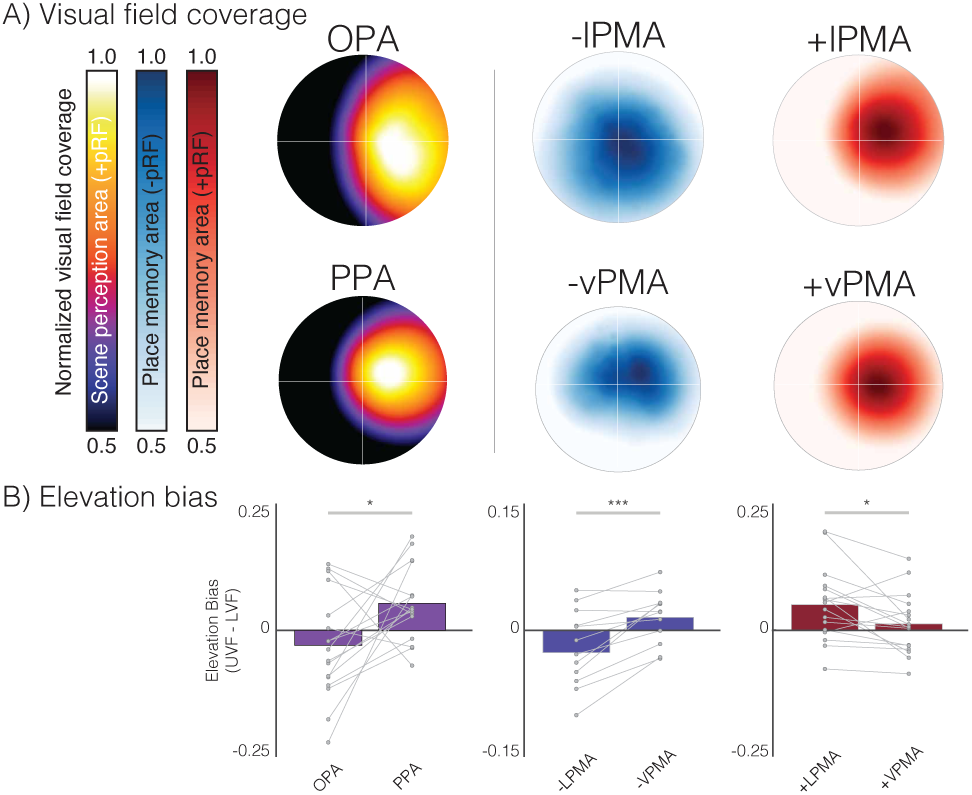
Shared visual field representations between paired perception and memory areas. **A.** Ventral and lateral SPAs/PMAs differentially represent the upper (ventral areas) and lower (lateral areas) visual fields. Group visual field coverage plots for SPAs and PMAs. X-axis represents ipsilateral to contralateral visual field (aligned across hemispheres), Y-axis represents elevation. All regions show the expected contralateral visual field bias. With respect to elevation, SPAs differentially represent the lower (OPA) and upper (PPA) visual field, respectively (left column). The same differential representation of the lower and upper visual fields is evident for -pRFs in LPMA & VPMA (middle column) but not for +pRFs in these areas (right column). The elevation biases are quantified in **B.** Bars represent average visual field coverage for contralateral upper versus lower visual field (UVF-LVF) with individual participant datapoints overlaid. *p<0.05, **p<0.01.

If, as hypothesized, +/− pRF populations reflect bottom-up sensory and top-down internal mnemonic processes, we predicted that the sign of the spatially-specific opponent interactions observed during visual mapping (Exp. 1) would reverse during a top-down memory paradigm. Specifically, Exp. 1 showed that when activity was high in +pRFs within the SPAs, activity was low in -pRFs within the PMAs. In contrast, we hypothesized that during a recall task, when -pRF activity would be high, +pRF activity would be low. To test for this competitive interaction during recall, in Experiment 2 we examined the activation profile of +/− pRFs during a place memory task, wherein participants recalled personally familiar visual environments (e.g., their kitchen, **Fig. 5A**). For this analysis, we first partitioned LPMA and VPMA into their +pRF and -pRF populations, respectively. Then, for each subject, we calculated the average activity (activation values versus baseline) from each of the 36 trials of the memory task for each population and ROI (e.g., -pRFs in the place memory areas, +pRFs scene perception area) and z-scored the activation values in each ROI. We compared the z-scored values using a partial correlation between the -pRF/+pRFs in the PMAs and the SPAs (e.g., correlation between -pRFs in LPMA with +pRFs in OPA, while controlling for +pRFs in LPMA, and vice versa) for each subject. Partial correlation allowed us to compare the unique impact of these distinct neural populations, while simultaneously controlling for non-specific effects (like motion and attention) that impact beta estimates on each trial. We compared the partial correlation values from the different pRF populations (i.e., -LPMA x OPA vs +LPMA x OPA) for all subjects using paired t-tests separately for the lateral and ventral surfaces.

**Figure 5.**
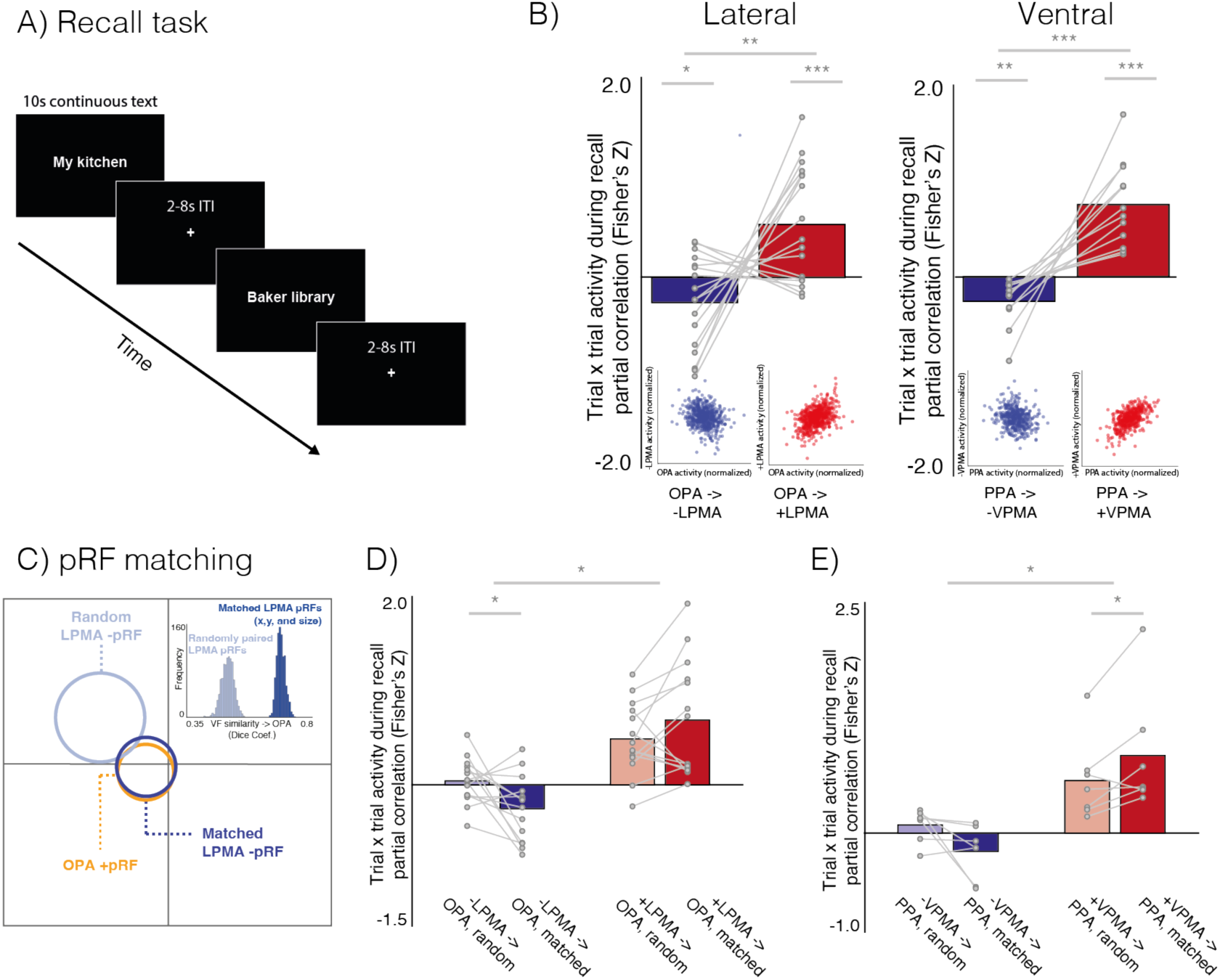
Positive pRFs in scene perception areas (SPAs) and negative pRFs in place memory areas (PMAs) exhibit a spatially-specific push-pull dynamic during memory recall. **A.** Recall task paradigm. In Exp. 2, participants visually recalled personally familiar places for 10 s (cued with stimulus name; 36 trials per participant), and we calculated activation versus baseline at each trial. **B.** Negative and positive pRF (-pRF, +pRF) populations in SPAs and PMAs exhibit an opponent interaction during recall. On both the lateral and ventral surfaces, when activity of -pRFs in the PMA is high, activity in the corresponding SPA is reduced. In contrast, when activity in +pRFs in a PMA is high, activity in the adjacent SPA is also high. Inset scatter plots show normalized trial-wise activation of each region for all trials in all participants. **C.** Schematic of pRF matching between OPA with -pRFs in LPMA. Schematic pRFs are depicted as circles. The OPA pRF is shown in orange, along with its best matched -pRF in LPMA (shown in dark blue) and randomly matched -pRF in LPMA (light blue). Inset. Histogram indicates the dice coefficients for iterative matching between OPA pRFs and randomly paired pRFs from LPMA (light blue) compared to matched -pRFs from LPMA (dark blue) for one example participant, showing better correspondence between matched vs random, “unmatched” pRFs. This pattern was consistent in all participants. **D,E**. Matched pRFs had a significantly stronger association in trial x trial activation than unmatched pRFs on the lateral **(D)** and ventral **(E)** surfaces, which suggests that the interaction between the perceptual and memory areas is spatially specific*: p < 0.05, **: p<0.01, ***: p<0.001.

As predicted, we observed an opponent relationship between activation of the -pRFs in the PMAs and the +pRFs in the SPAs during the place memory recall task (-LPMA x OPA vs +LPMA x OPA: t(16)=3.10, p=0.0006, D=1.52; -VPMA x PPA vs +VPMA x PPA: t(11)=5.27, p<0.0001, D=2.98). Specifically, on trials where the activity of -pRFs in PMAs was high, SPA activation was reduced **(Fig. 5B)** (t-test vs. zero – lateral t(16)=-2.19, p=0.04, D=0.53; ventral: t(11)=-3.63, p=0.003, D=1.04). In contrast, the +pRFs in the PMAs did not show a push-pull dynamic: on trials where activity of +pRFs in the PMA was high, activation of the SPA also increased (t-test vs. zero: lateral: t(16)=3.76, p=0.001, D=1.69); ventral: t(11)=5.88, p=0.0001, D=1.69) **(Fig. 5B)**. We found a similar pattern when we considered all trials from all subjects pooled together (**Fig. 5B insets, Fig. S4**), and we replicated this effect in an independently collected dataset **(Fig. S5)**.

Is the interaction between perceptual and mnemonic spatially specific? Importantly, if the interaction between the PMAs and SPAs is structured by a retinotopic code, we expect that the opponent dynamic between -pRFs in the PMAs with +pRFs in the SPAs would be stronger between pRFs representing shared regions of visual space. To test this, we again compared the partial correlation in trial x trial activation between +/− pRFs of the perception and memory areas, this time focusing on subpopulations of +/− pRFs with matched (versus unmatched) visual field representations (in x, y, and sigma) (see Methods; **Fig. 5C**). Because of our stringent pRF matching criteria (see Methods), the overall number of subjects included in this analysis was reduced (Lateral: N=14; Ventral: N=7).

This analysis revealed that the inhibitory interaction between -pRFs in PMAs and +pRFs in SPAs is structured by a retinotopic code (i.e., is spatially specific) **(Fig 5D, 5E)**. The opponent relationship between -pRF activation in the PMAs and matched +pRF activation in the SPAs was significant on both the lateral (F(1, 13)=16.45, p=0.001) and ventral (F(1, 6)=10.93, p=0.01) surfaces, reflecting the fact that –pRFs in PMAs exhibit a negative relationship with spatially-matched SPAs, whereas +pRFs in PMAs show a positive relationship with spatially-matched SPAs. Importantly, this relationship was modulated by matching on the lateral (Amplitude X Matching: F(1, 13)=4.64, p=0.05) and ventral (Amplitude X Matching: F(1, 6)=5.96, p=0.05) surfaces. Specifically, on the lateral surface, when trial x trial activity in SPA was high, activity in spatially-matched PMA -pRFs was relatively low (matched vs. zero: t(13)=2.73,p=0.01, D=0.75), and this anti-correlation was significantly stronger for matched vs. unmatched -pRFs (t(13)=2.48, p=0.02, D=1.01). On the ventral surface, a similar pattern was observed despite the small number of participants, but it did not reach significance (matched vs. zero: t(6)=1.83, p=0.11, D=0.76; matched vs. unmatched: t(6)=2.06, p=0.08, D=1.28). In contrast, for +pRFs in the PMAs, we observed a positive correlation with spatially-matched (vs. unmatched) +pRFs in the SPAs (Lateral matched vs. zero: t(13)=4.16, p=0.001, D=1.15; Ventral matched vs zero: t(6)=3.41, p=0.01, D=1.37). These data show that the spatially-specific opponent relationship between -pRFs in the PMAs and +pRFs in the SPAs evidenced during bottom up visual stimulation (i.e., pRF mapping in Exp. 1) is reversed during top-down memory recall, revealing a push-pull dynamic between +/− pRFs that represent similar regions of visual space.

Although the spatially-specific push-pull dynamic observed between +/− pRFs during both perception (Exp. 1) and recall (Exp. 2) is compelling, these two tasks represent situations wherein activation of the visual and memory systems should be maximally opposed (i.e., focusing attention externally on a visual stimulus vs. focusing attention internally during visual recall). Do -pRFs in the PMAs and +pRFs in the SPAs interact in a mutually inhibitory fashion in contexts where perceptual and memory systems are expected to interact, such as during familiar scene perception? As a final test of the interaction between the SPA and PMA pRFs, we asked whether the mutually inhibitory interaction between -pRFs in the PMAs and +pRFs in the SPAs is evident when participants view images of locations that were familiar to them in real life. We tested two possible hypotheses. On one hand, processing familiar scenes could be mutually excitatory for both the memory and perception areas, resulting in a positive trial x trial relationship between the SPAs and both +pRFs and -pRFs in PMAs. On the other hand, the opponent interaction might persist if the SPAs and -pRFs in the PMAs were interlocked in a mutually inhibitory relationship, like predictive coding^30^.

For this experiment (Exp. 3), a subset of participants (N=8) passively viewed portions of images (lower left and right quadrants) of recognizable Dartmouth College landmarks **(Fig. 6A).** We focused on the lateral surface (OPA and LPMA), because -pRFs could be localized most robustly in LPMA in our main sample of participants. Familiar scene stimuli were presented in the lower quadrants to maximally stimulate OPA and LPMA, and we specifically considered +/− pRFs with centers in the contralateral lower visual field. For this analysis, we tested whether the -pRFs in LPMA and +pRFs in OPA are correlated or anti-correlated during familiar scene perception. We hypothesized that activation of +pRFs in OPA would be inversely related to activation of -pRFs in LPMA, consistent with a mutually inhibitory interaction. Further, we predicted that this interaction would be spatially specific: the inhibitory dynamic should be stronger when visual information was presented in the contralateral lower visual field (i.e., preferred quadrant), relative to the ipsilateral lower visual field (i.e., non-preferred quadrant). Note that this is a coarser test of spatial specificity than that presented in Exp. 2 due to both the size of the stimuli presented in Exp. 3 and the comparison of preferred versus non-preferred quadrants (Exp. 3) as compared to matched versus unmatched individual pRFs (Exp. 2).

**Figure 6.**
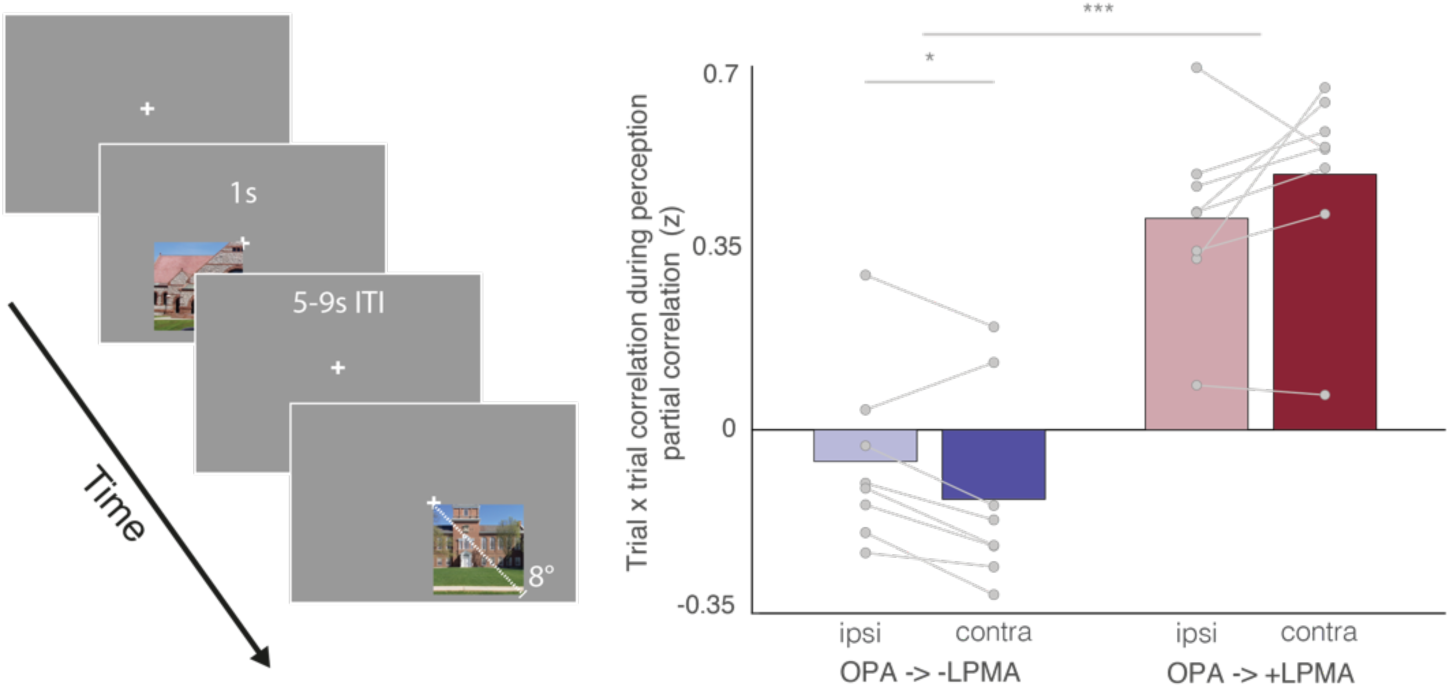
Positive pRFs in scene perception areas (SPAs) and negative pRFs in place memory areas (PMAs) exhibit a push-pull dynamic during perception of familiar scene images. **Left.** In Exp. 3, participants passively viewed portions of images depicting two familiar Dartmouth College landmarks in the lower visual field. We investigated the trial x trial correlation in activation between +pRFs in OPA and -/+ pRFs in LPMA. **Right.** - pRFs in LPMA exhibit an opponent interaction with +pRFs in OPA. 6/8 participants exhibited a negative correlation between these pRF populations, and this relationship was significantly stronger when stimuli were presented in the contralateral compared to ipsilateral visual field. In contrast, activation of the +pRFs in LPMA were significantly positive for both contralateral and ipsilateral visual field presentations. *: p < 0.05, ***: p<0.001.

We found that a spatially-specific, inhibitory interaction between +pRFs in OPA and-pRFs in LPMA persists during familiar scene viewing (Interaction between visual field and pRF association: F(1,7)=8.27, p=0.02; **Fig. 6, S7**). The trial x trial opponent dynamic between OPA pRFs, and -pRFs and +pRFs in the PMAs differed by visual field (t(7)=2.79, p=0.026, D=0.98). As we observed in recall, this dynamic was driven by the opposing sign of the relationship between pRFs in OPA with -/+ pRFs in LPMA. We found that the relationship between pRFs in OPA and -pRFs in LPMA was negative in 6/8 participants in the contralateral visual field, and the strength of the negative association between OPA and -pRF activity for stimuli in the contralateral visual field was significantly stronger than the ipsilateral visual field (t(7)=2.71, p=0.03, D=0.95). In contrast, the relationship between OPA and +pRFs in LPMA was positive in 8/8 participants, and was numerically stronger for the contralateral hemifield (t(7)=1.75, p=0.12, D=0.62). Together, the opposing responses during visual stimulation (Exp. 1 & 3) as well as during recall (Exp. 2), strongly suggest that the spatially-specific, mutually inhibitory interaction between -pRFs in mnemonic regions and +pRFs in perceptual regions is a generalizable description of their interaction that extends to naturalistic contexts, like familiar scene perception.

## Discussion

Here, we observe that a shared retinotopic code between externally-oriented (perceptual) and internally-oriented (mnemonic) areas of the brain structures their mutual activity, such that pRFs representing similar areas in visual space have highly anti-correlated activation during both bottom-up scene perception and also top-down memory recall. Together with recent reports showing retinotopic coding persisting as far as the ‘cortical apex’^15,16,31^, including the default mode network^15,16^, as well as the hippocampus^31,32^, our findings challenge conventional views of brain organization, which generally assume that retinotopic coding is replaced by abstract, amodal coding as information propagates through the visual hierarchy^7–10^ towards memory structures^11–14^. Furthermore, by examining the interaction between functionally paired perceptual and memory related areas, our work suggests that retinotopic coding may play an integral role in structuring information processing across the brain.

This conclusion has significant implications for our systems-level understanding of information processing at the cortical apex. Along with prior work in human and non-human primates^15,16^, our work demonstrates that high-level brain areas approaching the cortical apex ^12,13^ explicitly represent visual information in the environment. These regions have an inverted retinotopic code^5,17^ that organizes their functional interaction with visual areas. This view directly contrasts with the classic view of the cortical apex as an amodal and “internally-oriented” neural system. Large-scale deactivation of mnemonic areas at the cortical apex (e.g., the DMN) during visual tasks and activation during internally-focused tasks is among the most striking and widely-replicated network-level patterns in functional neuroimaging^33–36^. Critically, the deactivation within the DMN was not thought to represent stimulus information, but instead resulted from a trade-off between internally and externally oriented neural processes^35,36^. In contrast, our results suggest that the opponent activation profiles of DMN/non-DMN brain areas may reflect a mutually-suppressive functional interaction between mnemonic and perceptual areas that are involved in processing the same stimulus, rather than a trade-off between dissociable processes (like internal and external cognition). Thus, it may be necessary to reconsider the default mode network’s role in tasks that require externally-directed attention, as well as what representational content might be conveyed in negative BOLD responses.

How general is retinotopic coding for structuring perceptual-mnemonic interactions? On the one hand, because visually guided navigation has unique mnemonic demands (e.g., representing visual information out of view) scene areas might have privileged access to memory information^37^. Indeed, the scene memory and perception areas may uniquely span the boundary of DMN/visual cortex (**Fig. 1B, S3**)^23^ (but see^11^). Likewise, these unique mnemonic processes could lead to preferential coding of retinotopic information in scene memory areas, making the opponent interaction observed here “scene specific”. On the other hand, the present findings and others^15,31,32^ show that pRFs, including -pRFs, are widely distributed throughout the brain. As such, retinotopic representations could be well-poised to structure interactions between many functionally coupled brain areas broadly in cortex, beyond the domain of scenes, including other functional subnetworks situated within the default mode network^23,38–43^. Relatedly, matching visual (posterior) and language (anterior) areas have recently been identified at the anterior edge of visual cortex^11^, raising a question as to whether these areas might also be coupled via opponent dynamics that structure their communication. Beyond retinotopic codes, it is important to consider that the format of visual spatial coding within a region may depend on the computational roles a given region plays. For example, parietal, occipital, and temporal areas utilize multiple different visuospatial coding motifs^8,44–46^, including retinotopic^8^, spatiotopic^44,45,47^, and head/body-centered formats^50^. Thus, while our data clearly show the mutual retinotopic code structures the shared activity between paired perceptual and mnemonic areas, future studies will be necessary to further elaborate the nature of the visuospatial coding spanning internally- and externally-oriented networks in the brain.

Surprisingly, the PMAs contained a large proportion of +pRFs that, unlike the -pRFs, did not share the biased visual field representations of their SPA counterparts. This raises two questions. First, what regions contribute visual information to the memory areas’ +pRFs? While the present data offers no definitive answer, it is possible that they represent information arriving locally from other visual areas like the visual maps in the dorsal intraparietal sulcus^48^ or via longer range connections like the vertical occipital fasciculus^49^. Second, what do the +pRFs contribute to the PMAs’ functions? The robust activation of both +pRFs and -pRFs in the PMAs during memory recall suggests that these populations have related activity and differ primarily in their response to visual stimulation. Notably, the PMAs are defined based on a single response property (response during recall of places versus people), and functional regions may contain sub-regions with different functional, structural, and cytoarchitectonic characteristics (similar to the presence of multiple retinotopic maps within larger, functionally-defined visual regions like OPA^50^). Future work is needed to elucidate what differential roles the +/− pRFs in the PMAs might play.

To conclude, our results and others^15,16,31^ show that retinotopy is a coding principle that straddles internally-oriented (mnemonic) and externally-oriented (perceptual) areas in the brain. The inverted retinotopic code at the cortical apex is functionally tied to the positive retinotopic code in perceptual areas and may be crucial for scaffolding communication between memory and perception.

## Methods

### Participants

17 adults (13 female; age=22.8±3.5 STD years old) completed fMRI Experiments 1-2. A subset of 9 participants completed Experiment 3. Participants had normal or correct-to-normal vision, were not colorblind, and were free from neurological or psychiatric conditions. Written consent was obtained from all participants in accordance with the Declaration of Helsinki and with a protocol approved by the Dartmouth College Institutional Review Board (Protocol #31288).

### Visual Stimuli and Tasks

#### Population Receptive Field Mapping (Experiment 1)

During pRF mapping sessions, a bar aperture traversed gradually through the visual field, whilst revealing randomly selected scene fragments from 90 possible scenes. During each 36 s sweep, the aperture took 18 evenly spaced steps every 2 s (1 TR) to traverse the entire screen. Across the 18 aperture positions all 90 possible scene images were displayed once. A total of eight sweeps were made during each run (four orientations, two directions). Specifically, the bar aperture progressed in the following order for all six runs: Left to Right, Bottom Right to Top Left, Top to Bottom, Bottom Left to Top Right, Right to Left, Top Left to Bottom Right, Bottom to Top, and Top Right to Bottom Left). The bar stimuli covered a circular aperture (diameter = 11.4° of visual angle). Participants performed a color detection task at fixation, indicating via button press when the white fixation dot changed to red. Color fixation changes occurred semi-randomly, with approximately two-color changes per sweep. Stimuli were presented using PsychoPy (version 3.2.3) ^51^.

#### Scene Perception Area Localizer

The scene perception areas (SPAs, i.e. occipital place area, OPA; parahippocampal place area, PPA) are defined as regions that selectively activate when an individual perceives places (i.e., a kitchen) compared with other categories of visual stimuli (i.e., faces, objects, bodies) ^23,27,40,52^. To identify these areas in each individual, participants performed an independent functional localizer scan. On each run of the localizer (2 runs), participants passively viewed blocks of scene, face, and object images presented in rapid succession (500 ms stimulus, 500 ms ISI). Blocks were 24 s long, and each run comprised 12 blocks (4 blocks/condition). There was no interval between blocks.

#### Place Memory Area Localizer (Recall task; Experiment 2)

The place memory areas (PMAs) are defined as regions that selectively activate when an individual recalls personally familiar places (i.e., their kitchen) compared with personally familiar people (i.e., their mother)^23^. To identify these areas in each individual, participants performed an independent functional localizer. Prior to fMRI scanning, participants generated a list of 36 personally familiar people and places to establish individualized stimuli (72 stimuli total). These stimuli were generated based on the following instructions.

> “For your scan, you will be asked to visualize people and places that are personally familiar to you. So, we need you to provide these lists for us. For personally familiar people, please choose people that you know in real life (no celebrities) that you can visualize in great detail. You do not need to be contact with these people now, as long as you knew them personally and remember what they look like. So, you could choose a childhood friend even if you are no longer in touch with this person. Likewise, for personally familiar places, please list places that you have been to and can richly visualize. You should choose places that are personally relevant to you, so you should avoid choosing places that you have only been to one time. You should not choose famous places where you have never been. You can choose places that span your whole life, so you could do your current kitchen, as well as the kitchen from your childhood home.”

During fMRI scanning, participants recalled these people and places. On each trial, participants saw the name of a person or place and recalled them in as much detail as possible for the duration that the name appeared on the screen (10 s). Trials were separated by a variable ISI (4-8 s). Place memory areas were localized by contrasting activity when participants recalled personally familiar places compared with people (see ROI definitions section).

#### Perception of Familiar Scenes (Experiment 3)

A subset of participants in the main experiment (n=9) took part in this experiment, which involved viewing an image of two prominent Dartmouth College buildings (Baker Library and Rollins Chapel; one image per landmark). One participant was excluded for lack of familiarity with one building (Rollins Chapel). All remaining participants were familiar with the landmarks and had lived in the Hanover area for at least one year.

During scanning, participants passively viewed the familiar scene images. On each trial, participants maintained fixation while passively viewing the lower left or lower right quadrant of each image (display time: 1s). Images subtended as much of the whole lower visual field quadrant possible (0°-8° visual angle). We focused on the lower quadrants to maximally stimulate OPA and LPMA. Prior to scanning we showed participants the full image of each location to familiarize them with the specific image they would be seeing. Each image was presented eight times per location per run. Trials were separated by a variable ISI (4-8 s). We collected two imaging runs, resulting in 32 trials for each visual field location.

### fMRI Data Processing

#### MRI Acquisition

All data were collected at Dartmouth College on a Siemens Prisma 3T scanner (Siemens, Erlangen, Germany) equipped with a 32-Channel head coil. Images were transformed from dicom to nifti format using dcm2niix (v1.0. 20190902) ^53^.

#### T1 image

For registration purposes, a high-resolution T1-weighted magnetization-prepared rapid acquisition gradient echo (MPRAGE) imaging sequence was acquired (TR = 2300 ms, TE = 2.32 ms, inversion time = 933 ms, Flip angle = 8°, FOV = 256 × 256 mm, slices = 255, voxel size = 1 ×1 × 1 mm). T1 images segmented and surfaces were generated using Freesurfer ^54–56^ (version 6.0) and aligned to the fMRI data using align_epi_anat.py and @SUMA_AlignToExperiment^57^.

#### Functional MRI Acquisition

FMRI data were acquired using a multi-echo T2*-weighted sequence. The sequence parameters were: TR=2000 ms, TEs=[14.6, 32.84, 51.08], GRAPPA factor=2, Flip angle=70°, FOV=240 x 192 mm, Matrix size=90 x 72, slices=52, Multi-band factor=2, voxel size=2.7 mm isotropic. The initial two frames of data acquisition were discarded by the scanner to allow the signal to reach steady state.

#### Preprocessing

Multi-echo data processing was implemented based on the multi-echo preprocessing pipeline from afni_proc.py in AFNI^58^. Signal outliers in the data were attenuated (3dDespike^59^). Motion correction was calculated based on the second echo, and these alignment parameters were applied to all runs. The optimal combination of the three echoes was calculated, and the echoes were combined to form a single, optimally weighted time series (T2smap.py). Multi-echo ICA denoising^19,60–62^ was then performed (see *Multi-echo ICA*, below). Following denoising, signals were normalized to percent signal change.

##### Multi-echo ICA

The data were denoised using multi-echo ICA denoising (tedana.py^19,61,62^). In brief, PCA was applied, and thermal noise was removed using the Kundu decision tree method. Subsequently, data were decomposed using ICA, and the resulting components were classified as signal and noise based on the known properties of the T2* signal decay of the BOLD signal versus noise. Components classified as noise were discarded, and the remaining components were recombined to construct the optimally combined, denoised timeseries.

#### Population Receptive Field Modeling

Detailed description of the pRF model implemented in AFNI is provided elsewhere^18^. Briefly, given the position of the stimulus in the visual field at every time point, the model estimates the pRF parameters that yield the best fit to the data: pRF amplitude (positive, negative), pRF center location (x, y), and size (diameter of the pRF). Both Simplex and Powell optimization algorithms are used simultaneously to find the best time-series/parameter sets (amplitude, x, y, size) by minimizing the least-squares error of the predicted time-series with the acquired time-series for each voxel. Relevant to the present work, the amplitude measure refers to the signed (positive or negative) degree of linear scaling applied to the pRF model, which reflects the sign of the neural response to visual stimulation of its receptive field.

#### Sampling of fMRI Data to the Cortical Surface

For each participant, the analyzed functional data were projected onto surface reconstructions of each individual participant’s hemispheres in the SUMA standard mesh (std.141^63^), derived from the Freesurfer autorecon script using the Surface Mapping with AFNI (SUMA^57^) software and the *3dvol2surf* commands.

#### Regions of Interest (ROI) Definition

Scene perception areas (OPA, PPA) were established using the same criterion used in our prior work^23^. Scene perception areas were drawn based on a general linear test comparing the coefficients of the GLM during scene versus face blocks. Comparable results were observed when identifying the SPAs by comparing scene versus object blocks. A vertex-wise significance of p < 0.001 along with expected anatomical locations was used to define the regions of interest^52,64^.

To define category-selective memory areas, the familiar people/places memory data was modeled by fitting a gamma function of the trial duration for trials of each condition (people and places) using 3dDeconvolve. Estimated motion parameters were included as additional regressors of no-interest. Place memory areas were drawn based on a general linear test comparing coefficients of the GLM for people and place memory. A vertex-wise significance threshold of p < 0.001 was used to draw ROIs.

To control for differing ROI sizes across regions and people, we restricted all analyses to 300 vertices centered on the center of mass of each thresholded ROI. Consistent with prior work, we chose 300 vertices to ensure that no region or participant disproportionately contributed to any effects^23^. Results were qualitatively similar when 600 vertices were considered, suggesting our findings did not depend on ROI size.

#### ROI Analysis of pRF Amplitude

To calculate the percentage of -pRFs in each ROI we applied the following procedures. First, pRFs were thresholded on variance explained by the pRF model (R^2^>0.08), which is consistent with prior work using R^2^ thresholds ranging between 0.05 to 0.1^16,32,65,66^. Importantly, our results were consistent at R^2^ thresholds between 0.05 to 0.2. Next, to avoid analyzing only very few pRFs within an ROI, only ROIs consisting of >25 suprathreshold pRFs (>8.3 % of total ROI) were included. The percentage of suprathreshold pRFs with a negative amplitude within each ROI was then calculated and submitted for statistical analysis.

#### Visual Field Coverage

Visual field coverage plots represent the sensitivity of each ROI to different positions in the visual field. To compute these, individual participant visual field coverage plots were first derived. These plots combine the best Gaussian receptive field model for each suprathreshold voxel within each ROI. Here, a max operator is used, which stores, at each point in the visual field, the maximum value from all pRFs within the ROI. The resulting coverage plot thus represents the maximum envelope of sensitivity across the visual field. Individual participant visual field coverage plots were averaged across participants to create group-level coverage plots.

To compute the elevation biases we calculated the mean pRF value (defined as the mean value in a specific portion of the visual field coverage plot) in the contralateral upper visual (UVF) and contralateral lower visual field (LVF) and computed the difference (UVF-LVF) for each participant, ROI and amplitude (+/−) separately. A positive value thus represents an upper visual field bias, whereas a negative value represents a lower visual field bias. Analysis of the visual field biases considers pRF center location (like the center of mass calculation does), as well as pRF size and R2. This makes mean pRF value a preferable summary metric than analyzing pRF center position alone^18,32,40,50^.

#### Reliability of pRF Amplitude and Visual Field Coverage

To quantify the reliability of pRF estimates, we conducted a series of split-half analyses. First, pRF models were computed on the average time-series of all odd runs (1, 3, 5) and even runs (2, 4, 6), separately for each participant. Then, for each ROI, we identified all suprathreshold pRFs (R^2^>0.08) and pooled these pRFs across participants. This was done separately for +pRFs and - pRFs in each split (Odd, Even). Next, we conducted three tests of split-half reliability. First, we computed the percentage of pRFs in each ROI whose amplitude sign (i.e., positive or negative) remained consistent across splits. Second, to determine whether the sign of pRF amplitudes was reliable, we computed the Pearson’s correlation coefficient in amplitude across independent splits (Odd, Even), separately for positive and negative pRFs. We compared these R-values against zero (i.e. no correlation) using t-tests (two-tailed). Third, to determine whether the ROI visual field preferences are reliable, we computed visual field coverage maps from all suprathreshold pRFs in each ROI and split. Next, we calculated the dice-coefficient in the overlap between the split-half coverage maps and tested these values against zero (i.e. no overlap) using t-tests (two-tailed).

#### Recall Trial x Trial Analysis

To assess the interaction between +pRFs in SPAs and -pRFs in SPAs during memory recall, we adopted the following procedures. For each participant and ROI, we sampled the pattern of activity (t-value versus baseline) elicited during the recall of each personally familiar place (36 places per participant) from the place memory localizer experiment. Suprathreshold pRFs were separated according to amplitude (+pRFs, -pRFs) before averaging the recall responses across pRFs. This produced 36 responses (one for each recalled place) per pRF amplitude in each participant/ROI. We then z-scored these values for each participant and ROI separately.

After normalization, in each participant, we computed the partial correlation (Pearson’s R) between responses during memory recall of +pRFs in SPAs with -pRFs in PMAs, while controlling for the responses of +pRFs in PMAs and vice-versa. To determine whether the +pRFs and -pRsF from the PMAs had differential influence on activity in the SPAs, we compared the correlation coefficients from each population against each other using paired t-tests. To determine whether the influence of the -pRFs and +pRFs was significant, we compared the Fisher’s transformed correlation coefficients from each population (+pRFs and -pRFs in the PMAs) against zero (no correlation).

#### Spatial Specificity Analysis

First, we matched +pRFs in the scene perception areas with both +/− pRFs in the place memory areas separately (e.g., OPA -> -LPMA & OPA -> +LPMA) using the following procedure. On each iteration (1000 iterations total; randomized PMA pRF order): 1) For every pRF in a PMA, we computed the pairwise Euclidean distance (in x, y, and sigma) to all +pRFs in the paired SPA and found the SPA pRF with the smallest distance that was smaller than the median distance of all possible pRF pairs, 2) we required that all pRF matches were uniquely matched, so if an SPA pRF was the best match for two PMA pRFs, then the second PMA pRF was excluded. To prevent under sampling, only subjects with greater than 10 matched -/+ pRFs on average in each region were considered (Lateral surface: N=14, Ventral surface: N=7).

Second, we compared the correlation in trial x trial activation matched (versus non-matched) pairs of pRFs with “unmatched” pRFs. To create the “unmatched”, random pRF pairings, we randomly sampled pRFs in the memory area (repeated 1000 times). We then computed the unique correlation in trial x trial activation during recall between SPA pRFs and PMA pRFs, using the same procedure as in our main analysis (e.g., the partial correlation between SPA pRFs with PMA -pRFs, controlling for PMA +pRFs) for each iteration of the pRF matching. We compared the mean of the Fisher transformed partial correlation values across the iterations for the matched pRFs with the mean of random (i.e., unmatched) pRFs. To ensure that matched pRFs had better corresponding visual field representations than unmatched pRFs, we calculated the visual field overlap between pRF pairs in the matched samples, compared with the random samples (average dice coefficient of the visual field coverage for all matched versus unmatched iterations).

#### Perception of Familiar Scenes Trial x Trial Analysis

To assess the interaction between pRFs in OPA and +/− pRFs in LPMA during perception, we adopted the following procedure. We modeled task-evoked activity using a GLM with each image presentation fit as a separate regressor and calculated the average trial-wise activation of pRFs in OPA and +/− pRFs in LPMA. Importantly, because of our interest in spatially-specific interactions, we only considered pRFs with centers in the contralateral lower visual field, where the stimuli were presented (i.e., left hemisphere OPA pRFs had to have centers in the lower right quadrant). We then Fisher-transformed these values for each participant and ROI separately.

After normalization, in each participant,we calculated the partial correlation between the trial x trial activation of OPA with the negative and positive pRFs in LPMA separately for each hemifield (i.e., OPA with -pRFs in LPMA, controlling for +pRFs in LPMA when images were presented in the left hemifield). We compared the contralateral preference in the association (Fisher transformed partial correlation values) between the +/− pRFs using a repeated measures ANOVA with visual field (ipsilateral, contralateral) and pRF association (OPA x +LPMA, OPA x -LPMA) as factors.

#### Statistical Analysis

Statistics were calculated using the R Studio package (version 1.3)^67^. We conducted repeated measures analysis of variance (rmANOVA) using the ezANOVA function from the ‘ez’ package^68^. Alpha level of p < 0.05 was used to assess significance. We applied Bonferroni correction for multiple comparisons where appropriate.

## Author contributions

AS, EHS, CER conceived of and designed the experiment. EHS and BDG contributed stimulus code. AS and BDG collected the data. AS processed the data. AS and EHS analyzed the data. AS, EHS, CER wrote and edited the manuscript.

## Data availability statement

Data will be made available via datalad upon publication.

## Code availability statement

Code used for data analysis will be made available via GitHub upon publication. Analysis code will be made available to editors and reviewers during review via Dropbox upon request.

## Acknowledgements

We thank Iris Groen for assistance with specific analysis code. This work was supported by the National Institute of Mental Health under award number 1R01MH130529-01 (C.E.R.). A.S. was supported by the Neukom Institute for Computational Science (A.S.) and E.H.S by the Biotechnology and Biological Sciences Research Council award number BB/V003917/1.

## Supplemental Material

### Individual participant amplitude maps

**Figure S1.**
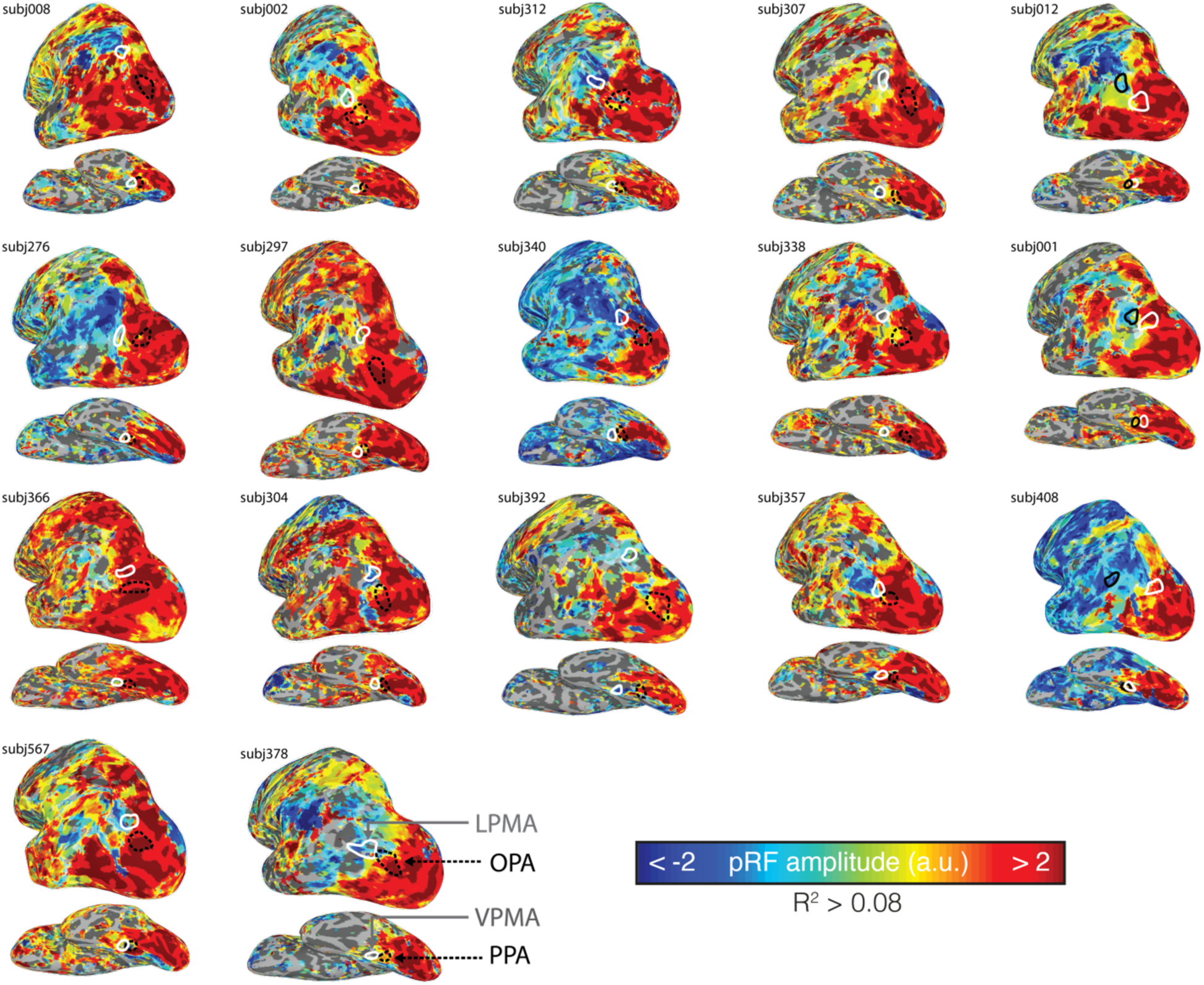
Transition from positive to negative amplitude population receptive fields (+pRF, -pRF) moving anteriorly from posterior cerebral cortex is evident in individual participants. Figure depicts amplitude maps from all participants’ left hemispheres. Only vertices surviving the threshold applied in the main text (R^2^ > 0.08) are shown. Individual participant SPAs and PMAs used for analysis are drawn in white (PMAs) and black (SPAs).

### Prominence of retinotopic coding within SPAs and PMAs

**Figure S2.**
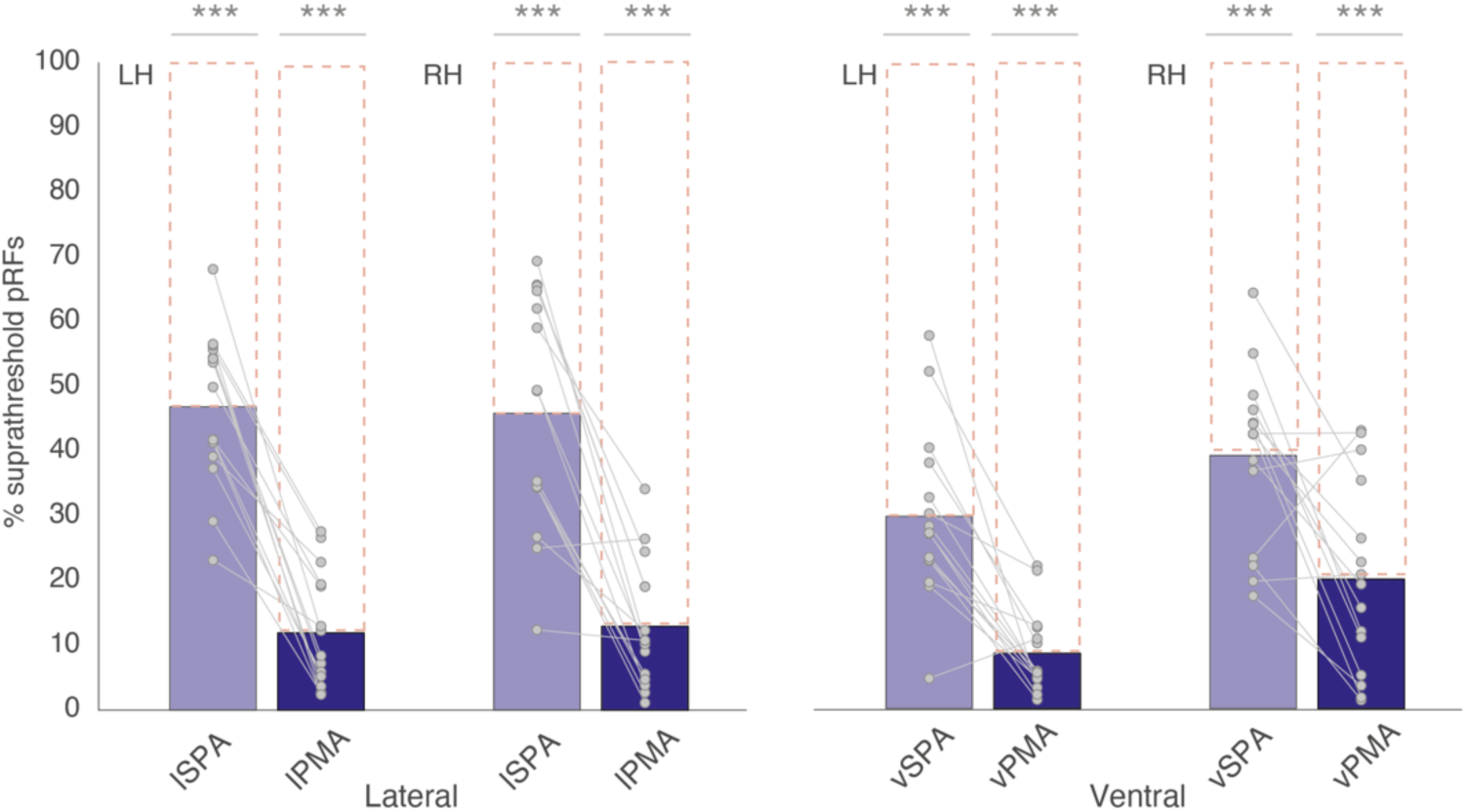
Retinotopic coding in SPAs and PMAs. To quantify the extent to which retinotopic coding is expressed within each ROI we first calculated the percentage of suprathreshold pRFs (R^2^ > 0.08) within our ROIs for each subject separately before testing each against a non-retinotopic prediction using t-tests (i.e., t-test versus zero, with Bonferroni correction). Retinotopic coding was significantly present within each ROI (LH; OPA: t(12.51)=, pcorr=3.35-9, D=2.99; PPA: t(16)=8.68, pcorr=5.67-7, D=2.17; LPMA: t(16)=6.23, pcorr=3.58-5, D=1.59; VPMA: t(16)=6.65, pcorr=1.64-5, D=1.66; RH; OPA: t(16)=12.15, pcorr=5.10-9, D=3.03; PPA: t(16)=11.97, pcorr=6.32-9, D=3.12; LPMA: t(16)=8.75, pcorr=5.05-7, D=2.18; VPMA: t(16)=6.39, pcorr=2.68-5, D=1.55). Bars represent the mean percentage of suprathreshold pRFs (R^2^>0.08) in each ROI/hemisphere for the lateral (left) and ventral (right) surfaces, respectively. Individual data points are overlaid. Each ROI exhibited a significant percentage of suprathreshold pRFs, ***p<0.001.

### Comparison of individual participant PMAs and SPAs to the default mode network

**Figure S3.**
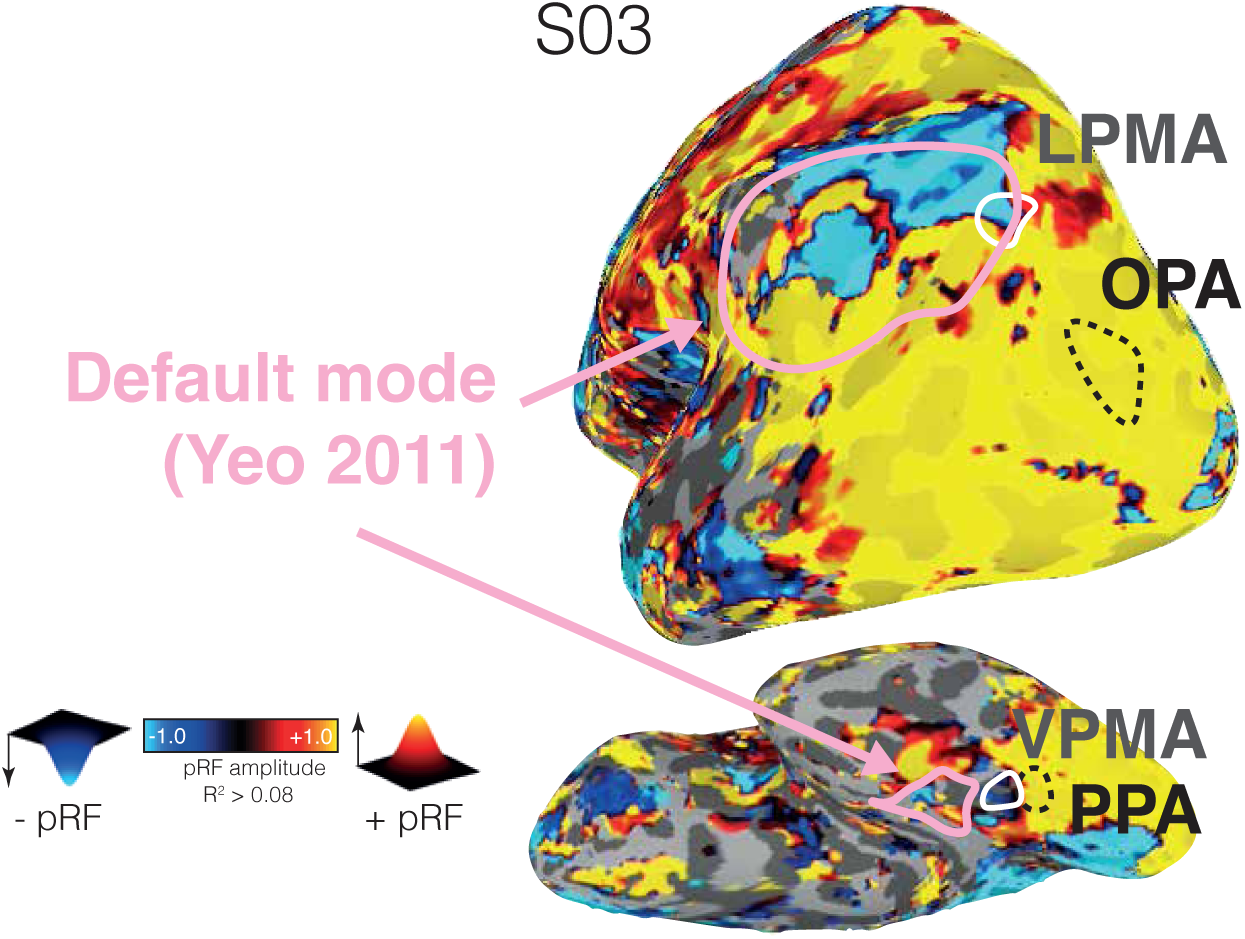
Comparison between the location of the SPAs, PMAs, and default mode network in one participant (example participant from Main text Fig. 2). This pattern was consistent in all individuals and at the group level (**Main text** Fig. 1B). Default mode network defined using the Yeo et al., 2011 parcellation.

### Differential correlation in trial x trial activation during recall across all trials

**Figure S4.**
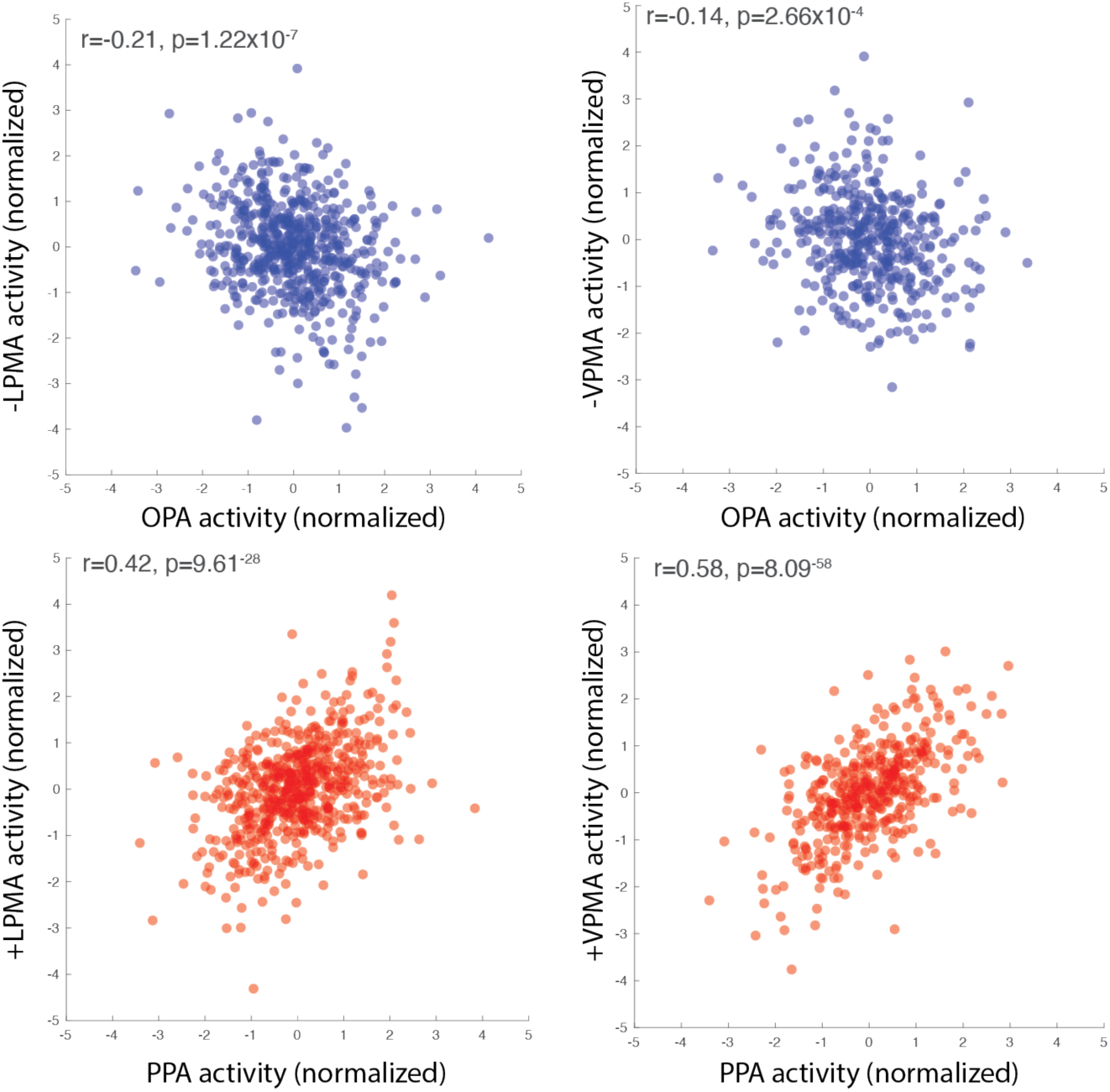
Experiment 2. Differential interaction between pRFs in SPAs with -/+ pRFs in memory areas is evident across all trials. Each scatter plot and corresponding correlation values depict the unique correlation between pRFs in the SPAs with -pRFs (blue) and +pRFs (red) in the PMAs (e.g., correlation between +pRFs in OPA with -pRFs in LPMA, controlling for +pRFs in LPMA). Each data point represents the z-scored activation on a given trial for all pRFS in the population (i.e., all -LPMA pRFS) for a given subject on a trial.

**Figure S5.**
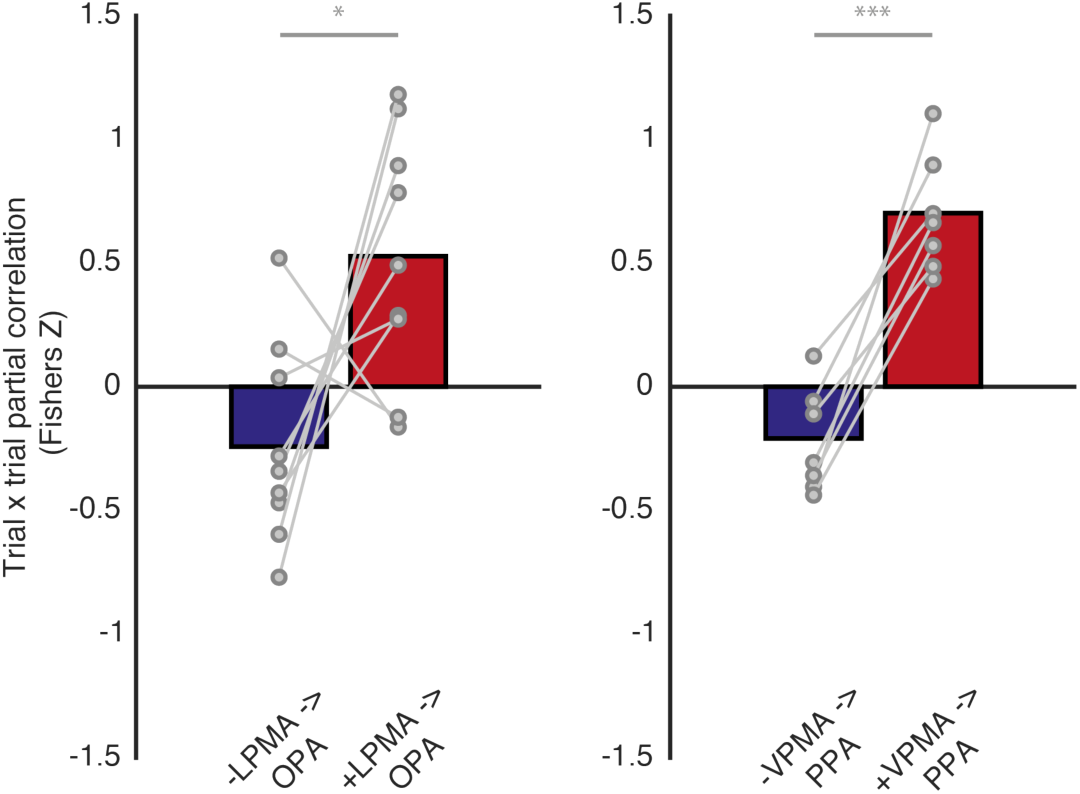
Trial x trial interaction between -/+ pRFs in the place memory areas and scene perception areas exhibit push-pull interaction in independent data. Recall trials were identical to the trials used in the localizer. Participants fixated on a dot projected in the center of the screen. They were then cued with the stimulus to be recalled for 1 second, followed by a 1 s dynamic mask, and 10 seconds of imagery. Trials were separated by a 4-8 s jittered interstimulus interval. Participants completed 32 imagery trials (16 for each landmark) separated into two imaging runs. One participant was excluded from the analysis for lack of familiarity with the landmarks; the remaining participants were familiar with the locations and had lived in the Hanover area for at least one year. Two participants did not have -pRFs in the ventral surface regions of interest. We tested for the relationship between +/− pRFs in the scene perception and place memory area using the same approach described in the Main text. We examined the unique correlation between the -/+ pRFs in the place memory areas and scene perception areas (i.e., correlation between activation of -pRFs in memory areas with pRFs in scene perception areas, while controlling for activation of +pRFs in the memory areas). We found evidence for the opponent interaction between -pRFs and +pRFs in this independent sample. We found that the relationship between the -/+ pRFs in the memory areas with the scene perception area pRFs was significantly different (Lateral – t(8)=2.61, p=0.018; Ventral – t(6)=7.82, p<0.0001). As we observed in our original analysis, the majority of participants showed a negative correlation in the trial x trial activation of the -pRFs in the place memory areas with pRFs in the scene perception areas (Ventral – 6/7 participants: t(6)= 2.79, p=0.031; Lateral – 6/8 participants: t(8)=1.79, p=0.11). Likewise, most participants showed a positive relationship between activation of +pRFs in the memory areas and pRFs in the perception areas (Ventral – 7/7 participants; t(6)=7.77, p=0.0002; Lateral – 7/8 participants; t(8)=3.30, p=0.01). This result gives us confidence that our original analysis was not influenced by potential circularity. * = p<0.05, *** = p < 0.005

**Figure S6.**
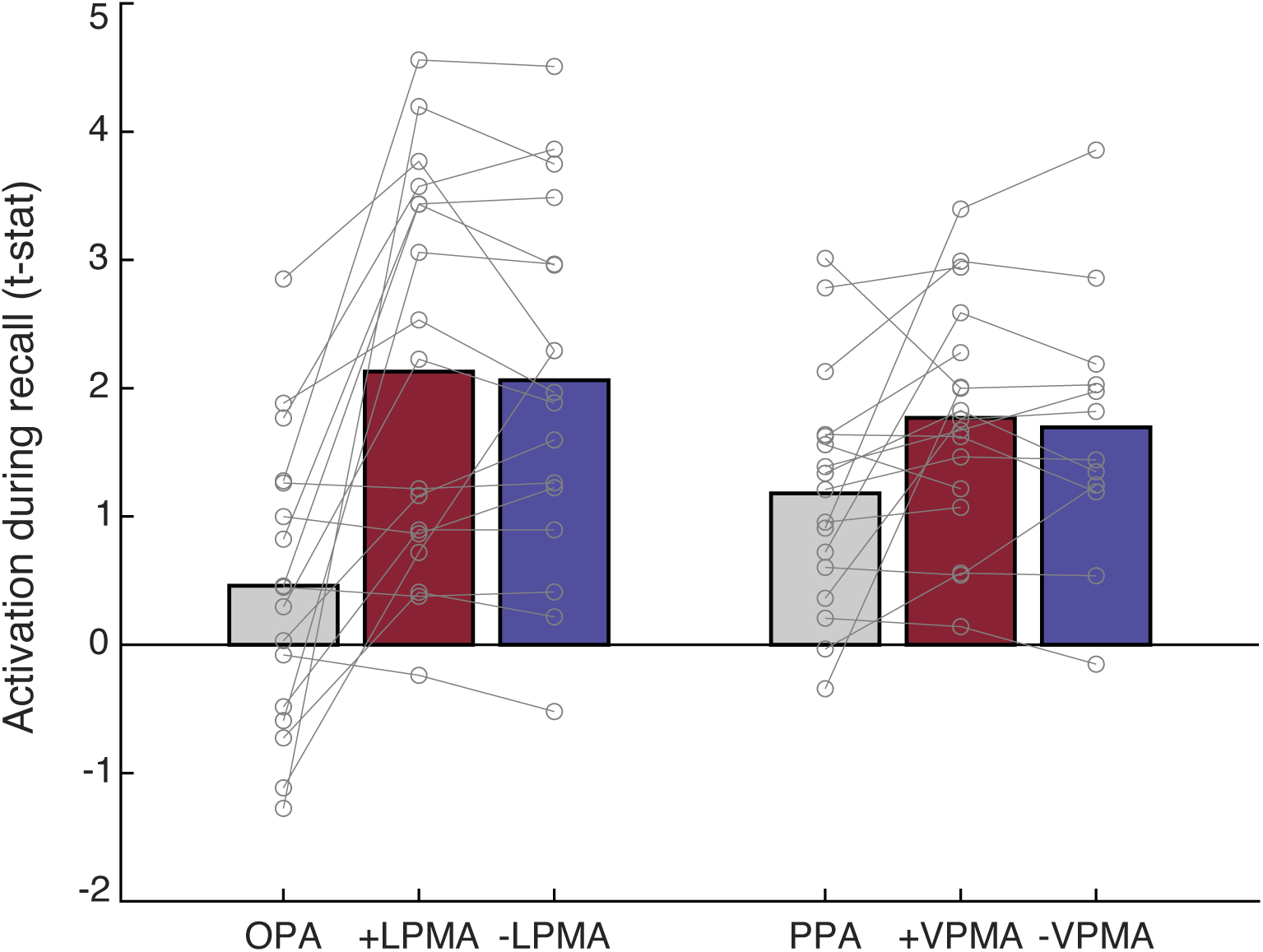
Experiment 2. Mean BOLD response amplitude relative to baseline during place recall trials for each ROI (OPA, LMPA) and pRF population (+/−).

**Figure S7.**
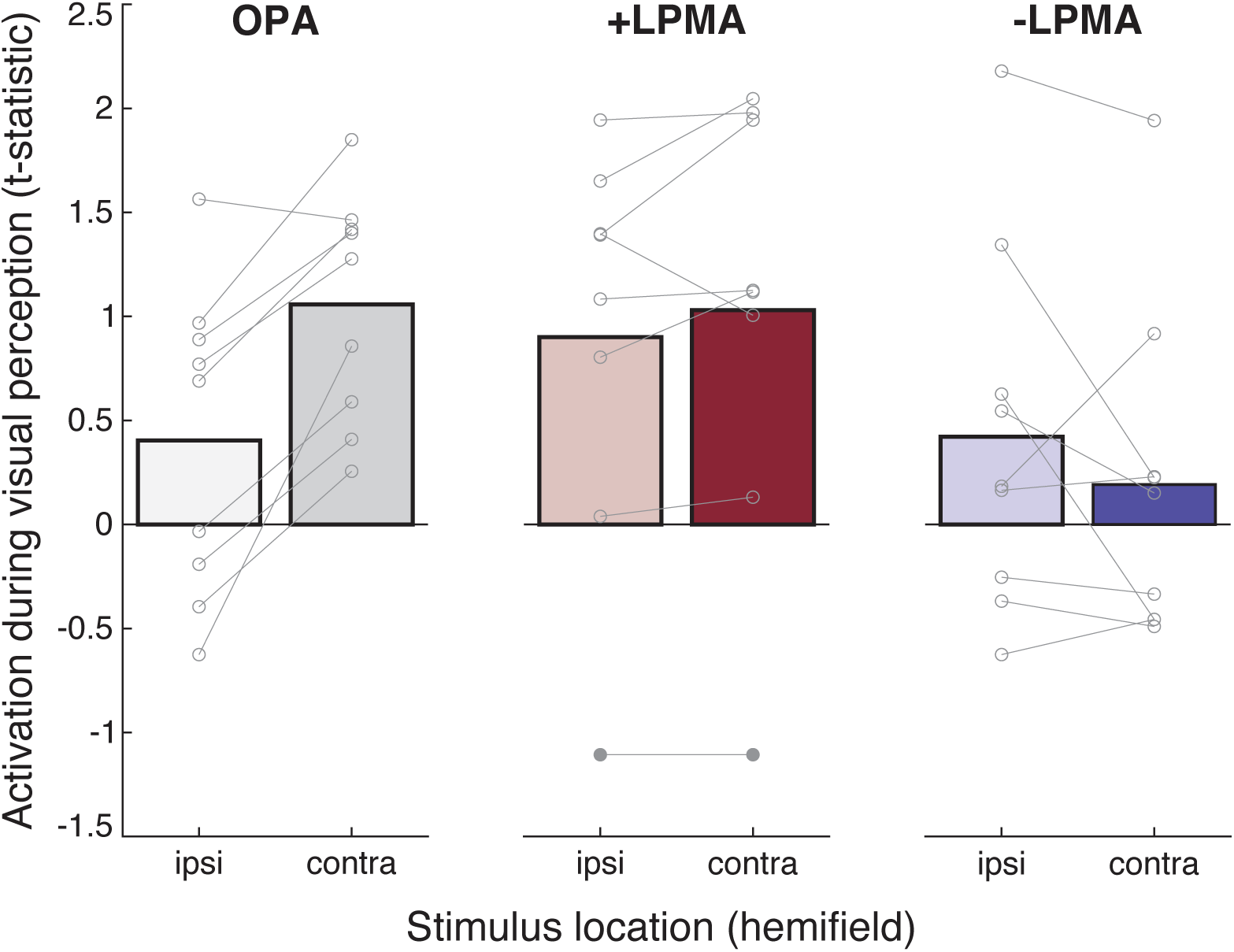
Experiment 3. Mean BOLD response amplitude relative to baseline when familiar scenes were presented in each lower quadrant (hemifield: ipsilateral and contralateral) for each ROI (OPA, LPMA) and pRF population (+/−).

